# Melanophore and fluoroleucophore synergistically photo-protect the Arabian killifish, *Aphanius dispar*, embryo from ultraviolet light

**DOI:** 10.1101/2024.11.30.626150

**Authors:** Maryam Alenize, Rashid Minhas, Tetsuhiro Kudoh

## Abstract

Pigment cells in fish species play crucial roles in forming colour patterns of each species and other physiological characteristics including photoprotection. Research on photoprotection by pigment cells in animals has primarily concentrated on black pigment cells, known as melanophores. However, the roles of other pigment cells and their synergistic effects on UV protection remain poorly understood. In this study, we use the Arabian killifish embryos as a model for studying the mechanisms of UV protection by different pigment cells. This species features highly fluorescent pigment cells called fluoroleucophores and black pigment cells known as melanophores. The fluorescent pigments and black melanin pigments are generated by genes *gch* (GTP cyclohydrolase) and *tyr* (tyrosinase) respectively. We generated *gch(−/−)* and *gch/tyr(−/−)* double mutant lines using CRISPR/Cas9 genome editing and examined the UV sensitivity of these mutant embryos. Both morphology and gene expression data revealed that the *gch/tyr(−/−)* double mutant line exhibited the highest UV sensitivity, and the *gch(−/−)* line also demonstrated a greater stress response compared to wild type (WT). From the study, we have identified the synergistic role of black and fluorescent pigment cells in providing effective UV protection from the early stages of embryonic development.

## Introduction

Pigment cells in fish play a crucial role in determining the colour and structure of their skin, scales, and other body parts, with speciation closely linked to these colour patterns. Additionally, colour patterns can vary significantly within closely related taxa, suggesting rapid evolutionary changes^1^. These diverse skin colour patterns serve multiple functions, including camouflage for predator avoidance, social signalling, mate evaluation and photo-protection^2,3^.

In fish, colours and patterns depend on various types of pigment cells, known as chromatophores^4^. Chromatophores originate from neural crest cells located at the dorsal neuroectodermal ridge, from which precursor cells migrate to form different tissues and structures^5,6^. Thus, the pigment patterns in fish are determined by their chromatophores’ types and arrangements, namely when and where they differentiate, migrate, and proliferate^7,8^.

Fish pigment cells interact with light in two primary ways, depending on the types of chromatophores involved. Some chromatophores, like melanophores, xanthophores, and erythrophores, absorb light, while others, such as iridophores and leucophores, reflect light.^9,10^. In teleosts, melanophores, iridophores, and xanthophores are widely distributed in many fish species, whereas leucophores are relatively rare^6,9,11,12^. Chromatophores contain intracellular organelles with pigments of varying biochemical compositions. For instance, melanophores contain melanosomes, leucophores have leucosomes, and iridophores possess reflecting platelets, each contributing specific colors and reflective patterns to the cells.

Melanophores are dendritic cells that align parallel to the skin’s layers and contain pigmented organelles known as melanosomes. These melanosomes house melanin, the dark pigment responsible for the brown or black coloration in fish. Their primary function is to disperse or aggregate melanin within the cytoplasm, enabling the fish to change color and providing protection from UV radiation^13^. Melanophores exhibit high sensitivity to light, allowing them to expand or contract in response

Leucophores, on the other hand, are white pigment cells that enhance the fish’s overall colour and brightness by scattering light with reflective platelets or crystals. Initially, leucophores contributed to the organism’s visual appearance, but they also play a crucial crucial role in camouflage, creating a silvery, reflective colouration that helps fish blend into their surroundings, especially in open water or variable lighting conditions.

Leucophores contain two types of pigment molecules: a weakly fluorescent pteridine pigment and a white pigment composed of urea^9,14^. Their reflective properties may also assist fish in maintaining an optimal body temperature by reflecting sunlight^6,9,11,12^. Generally, leucophores reflect white light of all wavelengths, while iridophores create iridescent colours^15^.

Fluoroleucophores, found in the Arabian killifish, are closely related to leucophores but exhibit greater fluorescence than those reported in other teleost fish species^16^.These cells have a similar dendritic morphology to leucophores and xanthophores ^15,16^. We have previously found GTP cyclohydrolase (*gch*) morphant and crispant show loss of the fluorescent pigment from the Arabian killifish indicating the crucial role of *gch* in generating the fluorescent pigment related to the group of pteridine molecules^16^.

Arabian killifish, *Aphanius dispar*, inhabits both freshwater and marine habitats and is widely distributed across the Middle East, from Western India and the Arabian Peninsula to West Africa. This species is commonly found in estuaries, lakes, and coastal areas, often residing in shallow waters, streams, and rocky habitats where it spawns in rock crevices ^17,18^. *A. dispar* is euryhaline, capable of tolerating a wide range of salinities, from freshwater to brackish and seawater^19^. According to Akbarzadeh *et al*., 2014, *A. dispar* can endure elevated temperatures, surviving up to 40 °C. Arabian killifish lives in the shallow water in the Middle East river and the sea, where plantation is highly limited, and embryos and fish are both exposed to strong sunlight. The presence of pigment cells developing from early stage of embryonic stage soon after the end of gastrulation^16,20^ suggests the importance of such pigment cells for UV protection in the habitats.

The sun is known to be the primary source of all forms of ultraviolet (UV) rays reaching the Earth’s surface. Ultraviolet radiation is divided into three major types based on the wavelength range measured in nanometers (nm): ultraviolet A (320-400 nm) has the longest wavelength with relatively harmless and moderate energy, Ultraviolet B (280-320 nm), with moderate harmful and high effective, and ultraviolet C the shortest wavelength (200-280) with severe dangers^21,22^. All these UV lights can cause DNA damage and oxidative stress of proteins and lipids in the cell with different extent of severity. UVA consists of up to 95% of the ultraviolet that reaches the earth’s surface, and it can penetrate the water column. Whereas UVB and UVC are filtered and absorbed by ozone, oxygen, and water within the atmospheric layer. However, because of ozone depletion, the earth’s surface has been exposed to an increased level of UVB and UVC^23,24^.

Fish embryos are particularly sensitive to UV light; exposure to UV radiation showed increased embryo malformations associated with decreased survival among exposed embryos^25–28^. Though there are many pigment cell types in the fish species, our knowledge in synergistic roles of these cells in UV protection is very limited. We have previously observed that in the Arabian killifish embryos, melanophore, fluoroleucophores and iridophores are neatly piled up as a unit with layer structure having iridophore at the skin surface, leucophore middle and melanophore in the deep layer suggesting synergistic role of these pigment cells and associated pigments in photoprotection^16^. In this report, we utilised CRISPR/Cas9 genome editing in Arabian killifish embryos to generate two specific pigment mutant lines: a *gch* mutant defective in fluorescent pteridine pigment synthesis in the fluoroleucophore and a *gch/tyr* double mutant which is devoid of melanin synthesis in the melanophores besides the *gch* mutation. We quantitatively analysed UV toxicity and stress response using morphological and gene expression analyses by qPCR. Our data indicates the synergistic role of melanophore and fluoroleucophore in UV protection, and physiological role of multiple pigment cells for the survival of fish in environments with intense sunlight.

## Materials and Methods

### Arabian killifish husbandry and embryos collection

The Arabian killifish *Aphanius dispar* wild type, *gch*^-^/^-^ and *gch−/−/tyr−/−* lines were sourced and maintained in the Aquatic Resource Centre at the University of Exeter. These fish were kept in a recirculation system under artificial photoperiods of 14 hours of light to 10 hours of darkness. The Arabian killifish *A. dispar* eggs were collected in egg spawning chambers and stored in Petri plates at a temperature of 28°C and a salinity of 35 ppt.

### Genome editing of *gch* and *tyr* gene by CRISPR/Cas9

To generate *gch* mutant and *gch/tyr* double mutant, two each CRISPR RNAs (crRNAs) were designed from the cDNAs of these two genes (Genbank accession numbers, PQ588425 for *gch* and PQ588426 for *tyr* respectively). The designed crRNA sequences for these genes were: *gch*_crRNA1 GTCCCGCTTACCCGCTCTGG and *gch*_crRNA2 GAGGAACTGAATGGCCTTGG), *tyr*_crRNA1 CTGGTCTCAGATGAACCCAA and *tyr*_crRNA2 AGTGTGCACTGATAATCTGA. Firstly, the mixture of two *gch*_crRNAs at 50 ng/µl each, 100 ng/µl tracrRNA and 200 nM Cas9 nuclease (NEB) was injected into the one-cell stage WT eggs with approximately 1nl volume. Embryos which have completely lost fluorescence were selected (88%) and raised to adult (*gch^-^/^-^* F0 generation). The *gch* mutant F1 line was established from natural spawning. Eggs were collected from these F1 fish and injected with mixture of two *tyr*_crRNAs to generate *gch/tyr* double mutant. All embryos which have completely lost the black pigment were selected (83%) and raised to adult (*gch−/−/tyr −/−* F0).

### DNA extraction, PCR and genotyping

For confirming the mutation in the *gch* and *tyr* genes, F1 generation of *gch/tyr* double mutant embryos were collected. Each embryo was snap frozen in liquid nitrogen, and subsequently dissolved in 50ul of 50mM NaOH and homogenised using an electronic pestle, incubated at 95 °C for 10 minutes, next, cooled down the mixture to 16 °C. Then, 5ul of 1M of Tris hydrochloride (1M Tris–Hcl, pH = 8.0) was added.

The targeted genomic region of *gch* or *tyr* gene was amplified by using Promega PCR Master Mix Green with using following volumes: 1 µl of genomic DNA mixture, 1 µl each forward and reverse primers (25 μM):, 12.5 Promega PCR Master Mix Green and nuclease free water to 25μl PCR reaction. Primer sequences flanking *gch* targeted region were: *gch*_F (AAAAGTTGGAGAAACCGCCG), *gch*_R (GATGGTCTCATGGTAGCCCT), *tyr*_F (GAAGGAGATGGCTCACCT), *tyr*_R (GCTAATGAGGTTGGGATT). The amplification conditions were as follow: 95°C 1min, 35 cycles (95°C 30 sec, 58°C 1 min, 72°C 1 min). The final extension step was performed at 72°C for 5 minutes. The PCR product was purified by Promega PCR clean-up system and cloned into a pGEM-T Easy Vector for sequencing.

### UVC radiation exposure

UVC radiation was provided by a UV crosslinker (analytic jena US, UVP Cross linker, CL-3000) containing six LED-lamps at wavelength 254 nm operated at 250 V. Ten embryos at 4dpf were placed in a Petri dish containing 25 ml of 35 ppt Artificial seawater. UVC lamps were placed at 17 cm above the dish’s surface. Four conditions were set up for exposure: No UV exposure (control) 0 mJ/cm^2^, UV 25 mJ/cm^2^, UV 50 mJ/cm^2^ and UV 100 mJ/cm^2^. Moreover, two conditions, 6 mJ/c m^2^ and 12.5 mJ/cm^2^, were tested before applying the above-mentioned doses. After UV irradiation, all exposed and non-exposed embryos were transferred to an incubator at 28°C for 24 hours with 12h light and 12h dark.

### Mortality, survival, heat beat rates and malformation

The effects of UVC exposure on the morphology and development of embryos were examined at 5 days post fertilization (5 dpf) until the hatching, following 24 hours of UV exposure. Mortality and malformation were analysed daily, with immediate removal of any dead embryos to avoid contamination. The mortality was conducted by observing locomotion, heartbeat and blood circulation using a Nikon SMZ1500 and an Olympus SZX16 microscope. The heart-beat rates of treated and untreated UV of *A. dispar* were monitored at 24-, 48- and 96-hours post-UV exposure. The heartbeats of all 10 embryos were observed for 30 seconds, and then the heart rate per minute was calculated. Before manual counting, embryos were kept at room temperature for 10 minutes to allow the heartbeat to stabilize. The malformation rates for UV exposure *A. dispar* embryos were assessed through microscopic examination at 24 hours post UV exposure until the hatching day and imaging for the main body changes. Pigment cell aggregation was observed microscopically in the designated regions. Every experiment was performed independently in triplicate.

### Imaging

For imaging purposes, the embryos were immersed in gel canals by leaving 1% agarose (sigma, A9539) made of deionised water to set on 1.2 mm diameter glass tubes (27). photographs were taken using a Nikon DS-L3 (DS-Fi2-L3u) camera and NIS-Elements software (version 4.30) on Nikon SMZ1500 microscope using normal incident light and appropriate GFP and RFP filters.

### RNA extraction, cDNA synthesis and Real-time quantitative PCR (qPCR)

The total RNA was extracted from embryos after 24 hours post-UV exposure from both untreated control and treated UV groups. Five embryos from each of the four conditions: Control (untreated), UV 25 mJ/cm^2^, UV 50 mJ/cm^2^, and UV 100 mJ/cm^2^, were collected from three different strains: A. dispar WT, A. dispar gch^-^/^-^, and A. dispar gch−/−tyr−/−. The collected embryos were placed in a tube containing 200 μl of Trizol reagent (RTI, Sigma T9424) and homogenized using an electric pestle. Equal volume of ethanol (95-100%) was added, and the total RNA was then isolated using Direct-Zol TM RNA miniprep kit) as per manufacturer’s protocol. RNA was eluted in in nuclear-free water and concentration was assessed using Qubit and Nanodrop, while RNA quality and integrity was verified using Tape Station. Subsequently, the RNA samples were stored at −20 °C.

The cDNAs were synthesised using the Thermo Scientific™ RevertAid™ kit as per manufactures’ protocol. All RNA samples were normalised to equal concentration (200 ng/ul) before setting up the reaction. Each reaction contained 1ul of RNA, 1 μl of Random hexamer primer and nuclease-free water to achieve a total volume of 12 μl for each reaction. Subsequently, 5X Reaction Buffer (4 μl), RiboLock RNase Inhibitor (20 U/µL) (1 μl), 10 mM dNTP Mix (1 μl) and of RevertAid M-MuLV RT (200 U/µL) were added then centrifuge briefly. The reaction was then incubated at 42 °C for 60 minutes then terminated the reaction by heating at 70 °C for 5 minutes. cDNA samples were stored at −20 °C or −80 °C.

Real time PCR was performed using Luna Universal qPCR Master mix (NEB, Catalogue M3003S) in a total volume of 10 µl. Each reaction contained 5 µl of Luna mix Universal qPCR Master mix, 0.5 µl of forward and reverse primers (10 µM), 1 µl of cDNA 3 µl of NF water. qPCR was performed on QuantStudio™ 3 Real-Time PCR System with the following cycling conditions: initial denaturation at 95°C for 1 minutes, followed by 35 cycles of denaturation at 95 °C for 15 seconds, and extension at 60 °C for 30 second.

### Ethical declaration

All experiments and methods were performed in accordance with relevant guidelines and regulations of UK Home Office with the licence number PP4402521. All methods are reported in accordance with ARRIVE guidelines. All experiments and methods approved by the Animal Welfare and Ethical Review Board (AWERB) at the University of Exeter.

### Data availability

The datasets generated during the current study are available in the Genbank (accession numbers, PQ588425 for *gch* and PQ588426 for *tyr* respectively). [These data will be available upon publication.]

## Results

### Generation of fluorescent and black pigments mutants

To generate pigment mutant lines defective in fluorescent pigment (*gch* mutant) and both fluorescent and black pigments (*gch/tyr* double mutant), the gch gene was initially knocked out by injecting two crRNAs as mixture (*gch*_crRNA1 and *gch*_crRNA2). 88% of injected embryos showed complete loss of fluorescent pigment. These non-fluorescent embryos were raised to adult to establish the F0 founders of the *gch* mutant line. Eggs were then collected from these F0 fish through natural spawning and raised to form the F1 generation.

To generate *gch/tyr* double mutant, eggs were collected by natural spawning from the *gch* mutant F1 generation and injected with two *tyr*_crRNAs (*tyr*_crRNA1 and *tyr*_crRNA2). Among these injected embryos, 83% showed a complete loss of black pigment. These embryos were raised to adult to establish *gch/tyr* double mutant founders.

The adult *gch* mutant fish did not show any noticeable differences in colour pattern compared to wild-type fish (Fig. 1A. A’, B, B’). However, the *gch/tyr* mutant fish displayed a clear loss of dark pigments and showed typical albino colour pattern in both the skin and lens (Fig. 1A, A’ and C, C’).

**Figure 1.**
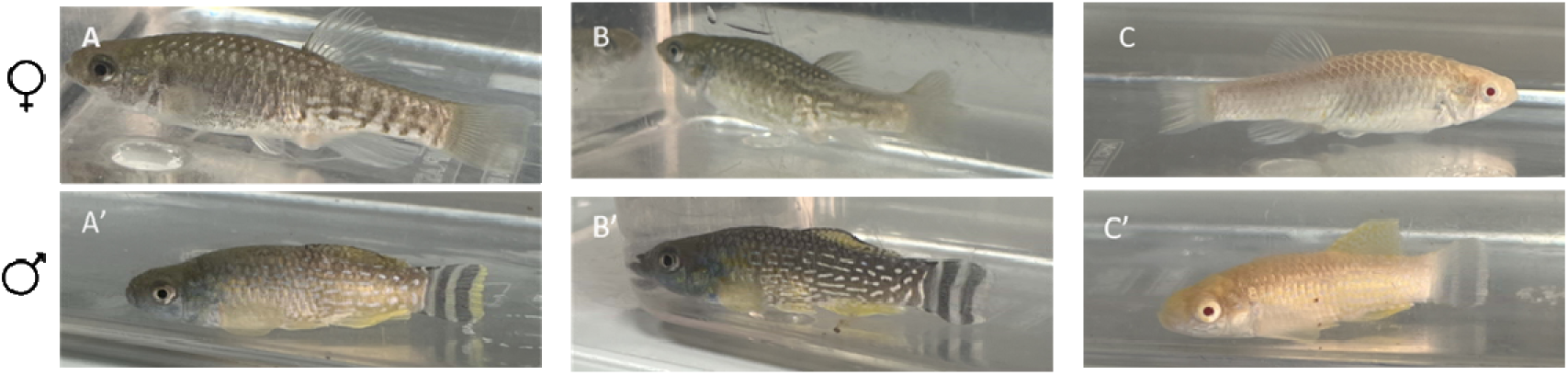
Arabian killifish *Aphanius dispar* adult fish with WT (A, A’), *gch* mutant (B, B’) and *gch/tyr* double mutant (C, C’) with females (A, B, C) and males (A’, B’, C’).

At the embryonic stage WT, *gch* and *gch/tyr* double mutants exhibited distinct and clear differences their colour pattern. As reported from the morphant and crispants from Hamied et al. 2020 ^16^ mutations in the *gch* caused loss of fluorescence from the fluoroleucophores (Fig. 2B, B’). In addition, with the mutations of *gch/tyr*, both fluorescence and black pigments were lost (Fig. 2C, C’).

**Figure 2.**
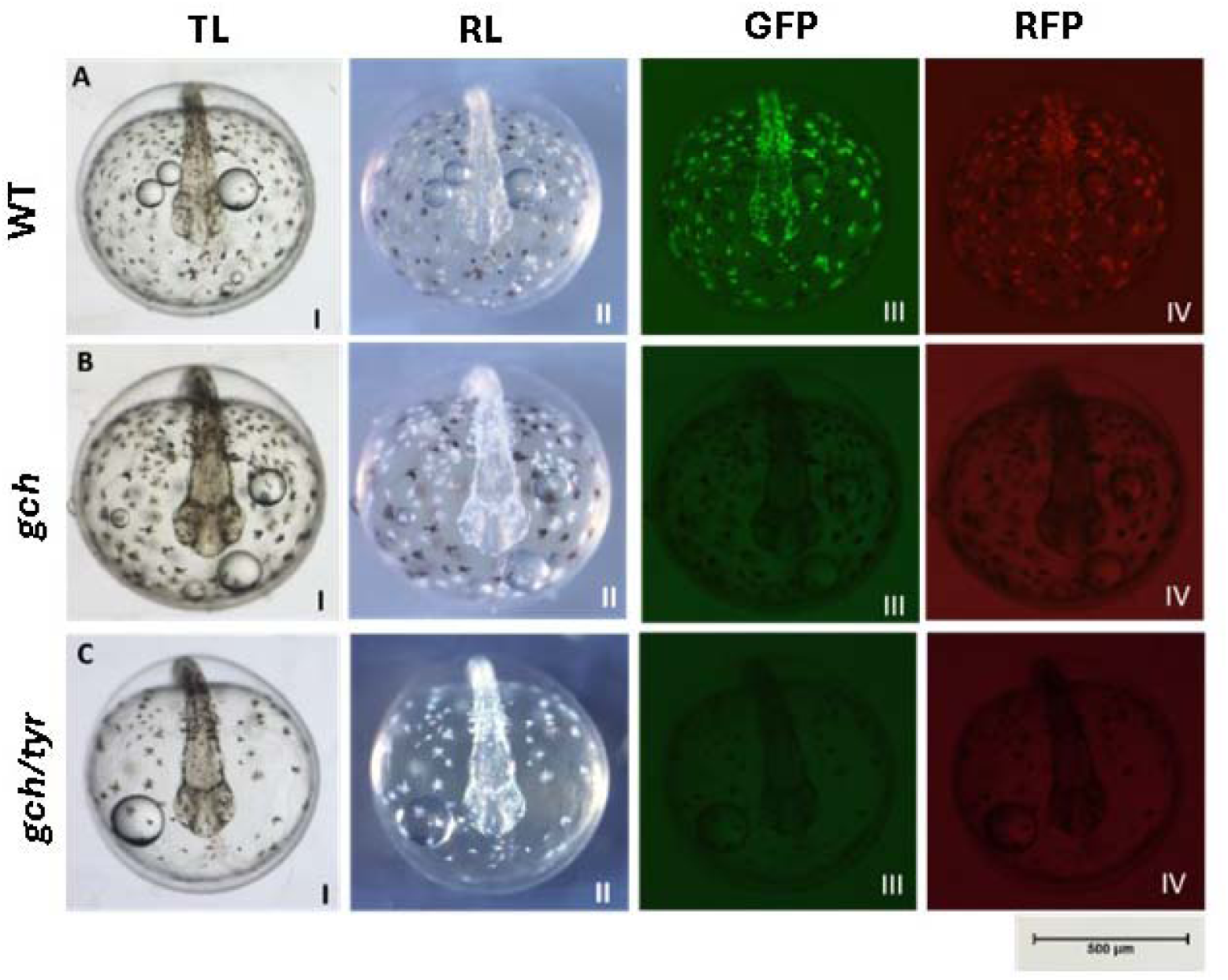
Arabian killifish *Aphanius dispar* whole e*m*bryos at 3 days post fertilization from anterior dorsal view. **A.** *A. dispar* wild-type WT, **B**. *A*. *dispar gch−/−* (induce loss fluorescent in leucophores), **C.** *A. dispar gch(−/−)/tyr(−/−)* (Group image in Fig. S1). TL. Transmission light. RL. Reflection light. GFP, fluorescent light with GFP filter. RFP. Fluorescent light with RFP filter.

### Genotyping of fluorescent and black pigments mutants

To confirm that mutations had indeed occurred in the *gch* and *tyr* genes in these mutant fish, the area of exon-1 of these two genes were amplified by genomic PCR using individual embryos from the *gch/tyr* double mutant. The amplified DNA was subcloned into the pGMT Easy plasmid vector and sequenced. Sequenced DNA showed mutations occurred both in the *gch* and *tyr* genes in the expected region around the two crRNA sequence in the exon. The mutations found in the *gch* showed small deletions both in the crRNA1 and crRNA2 regions (Fig. 3A) whereas mutations found in the *tyr* showed large deletion stretching from crRNA1 to crRNA2 sequence (Fig. 3B). Though the size of mutations were highly different between *gch* and *tyr* genes, we have confirmed specific deletion occurred in the designed exon (exon-1) of these genes causing frame shift mutations that are consistent with our observation of the loss of fluorescence and black pigments respectively (Fig. 2).

**Figure 3.**
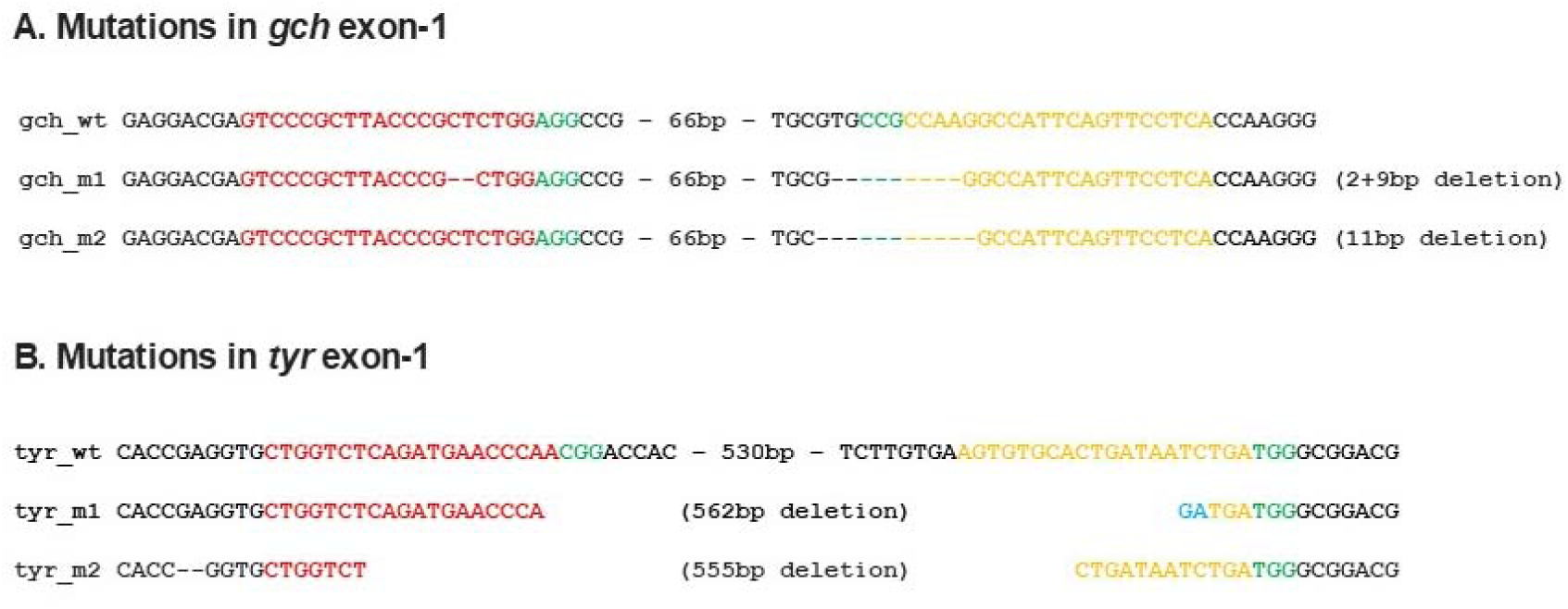
Mutations in the *gch* and *tyr* exon 1. **A.** and B, *gch* and *tyr* sequence in the exon-1 of each gene around the crRNA1 (red) and crRNA2 (orange) regions. PAM is highlighted in green.

### Impact of UV on mutant embryo viability

To investigate the effects of UV light we exposed WT, *gch* and *gch/tyr* mutant embryos to UV light. Embryos were collected from these strains through natural spawning, incubated for 4 days at 28 °C and then exposed to UVC at 4dpf with a variety of doses ranging from 25 to 100mJ/cm2 (Fig. 4). In embryos without UV exposure, melanophore and fluoroleucophore tend to maintain a distance of one to two cells diameters from each other (Fig. 4A, A’’, A’’’). However, in embryos exposed to UV, the shape of these pigment cells was more often distorted, and they were positioned much closer to each other (Fig.4B, C, D).

**Figure 4.**
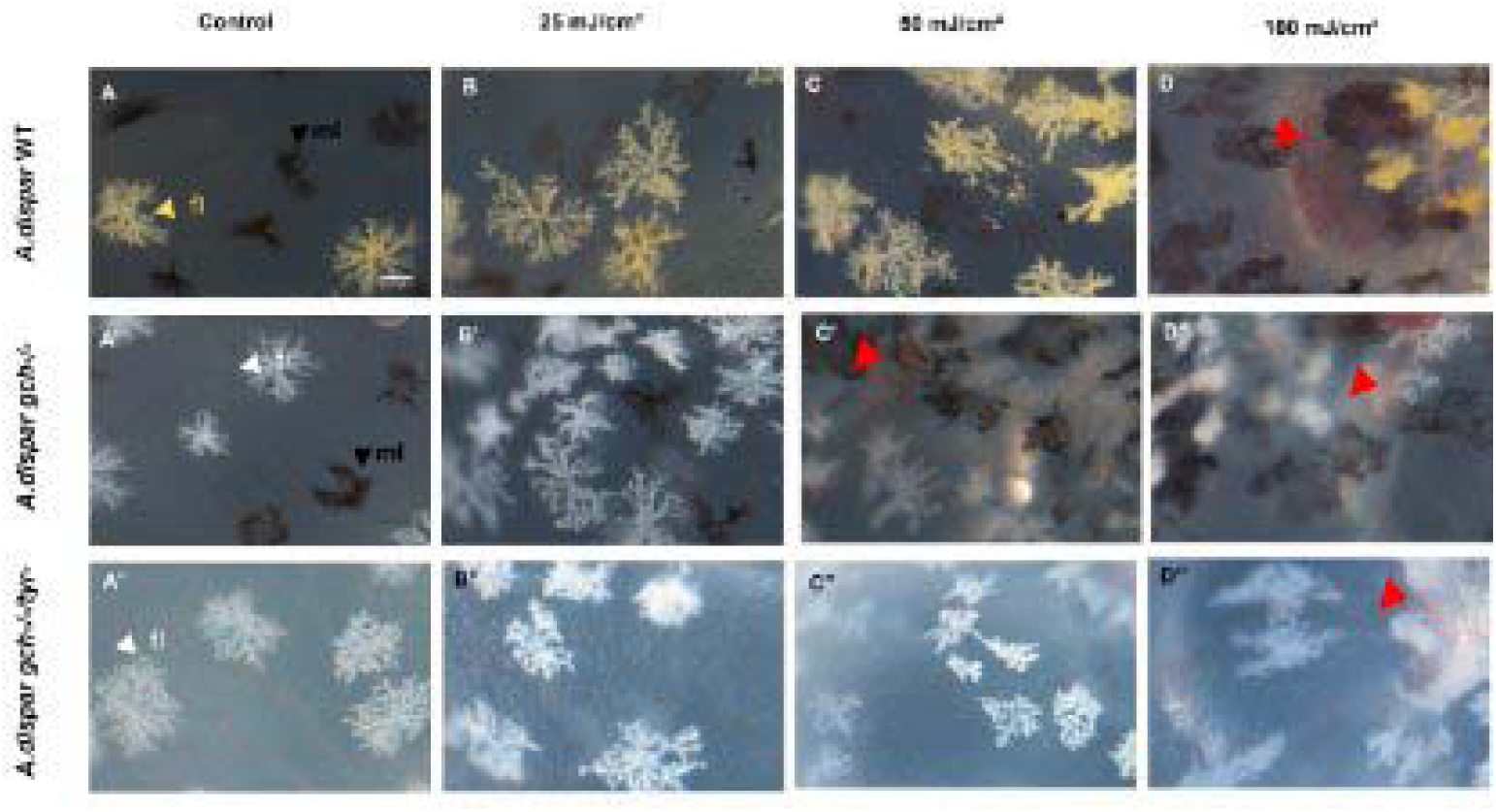
Live image of the *A. dispar* embryos after exposure to UV. The embryos were exposed to 25, 50 or 100mj/cm2 UVC at 4dpf (B, C,and D respectively), and imaged after 24h post exposure with focusing on fluoroleucophore (yellow or white) and melanophore (black) on the surface of the yolk. A-D, WT, A’-D’. *gch*, A’’-D’’ *gch/tyr*. Scale bar: 100um.

UV exposure also affected the survival of the exposed embryos in the following few days with dose dependent manner (Fig. 5). WT exhibited the highest survival rate even with even at the highest doses of 50 and 100mJ/cm2 (Fig. 5A). *gch* mutant showed slightly decreased survival whereas *gch/tyr* double mutant exhibited markedly reduced survival, even at lower doses (Fig. 4B, C).

**Figure 5.**
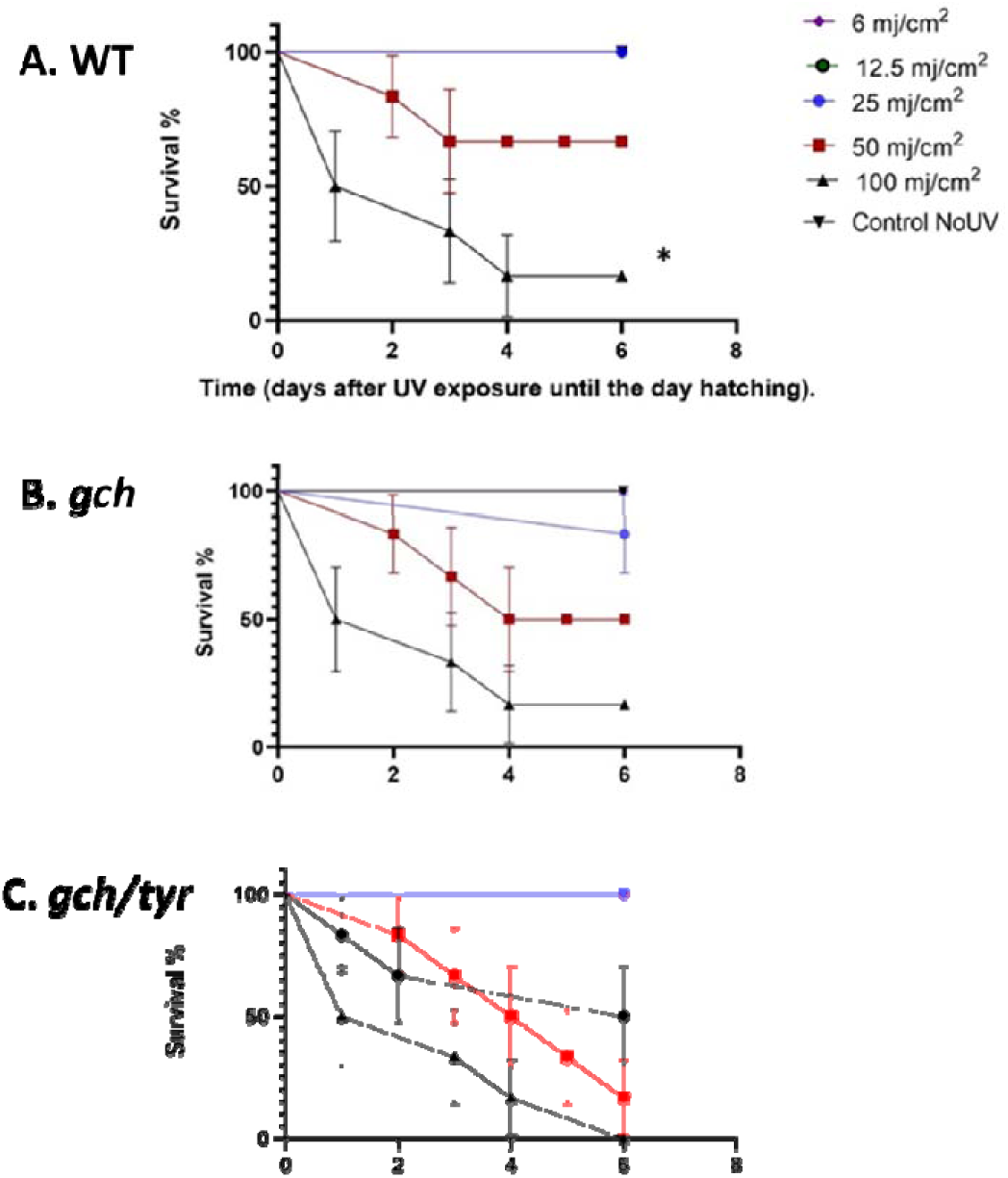
Survival of embryos after UV exposure. *A. dispar* embryos were exposed to UV at 4 dpf, and the survival was counted until 6 day post exposure. A. WT, B, *gch* and C. *gch/tyr* embryos. Overall WT shows the highest survival rates (A), while *gch* showed a slightly lower rate (B) and *gch/tyr* show severely reduced survival by UV exposure (C). Each point is the mean of three biological replicates ± SE. P-values between 0.05 and 0.001 were considered significant. * p < 0.05 and ** p < 0.01 compared to the control.

To examine additional milder phenotype of UV toxicity in the survived embryos from three genotypes, we have examined the heart rate of the UV exposed embryos at 24h and 48 post exposure (Fig. 6). UV exposure decreased heart rate across all genotypes in a dose-dependent manner. At 24hpe, WT embryos exhibited the highest resistance to the UV, maintaining higher heart rate (Fig. 6A) while *gch* and *gch/tyr* showed markedly reduced heart rate. In contrast, at 48 hpe, both WT and *gch* showed a similar dose dependent decline in heart rate, with *gch/tyr* having more severe decline, suggesting acceleration of the heat defect in the *gch* mutant compared to WT.

**Figure 6.**
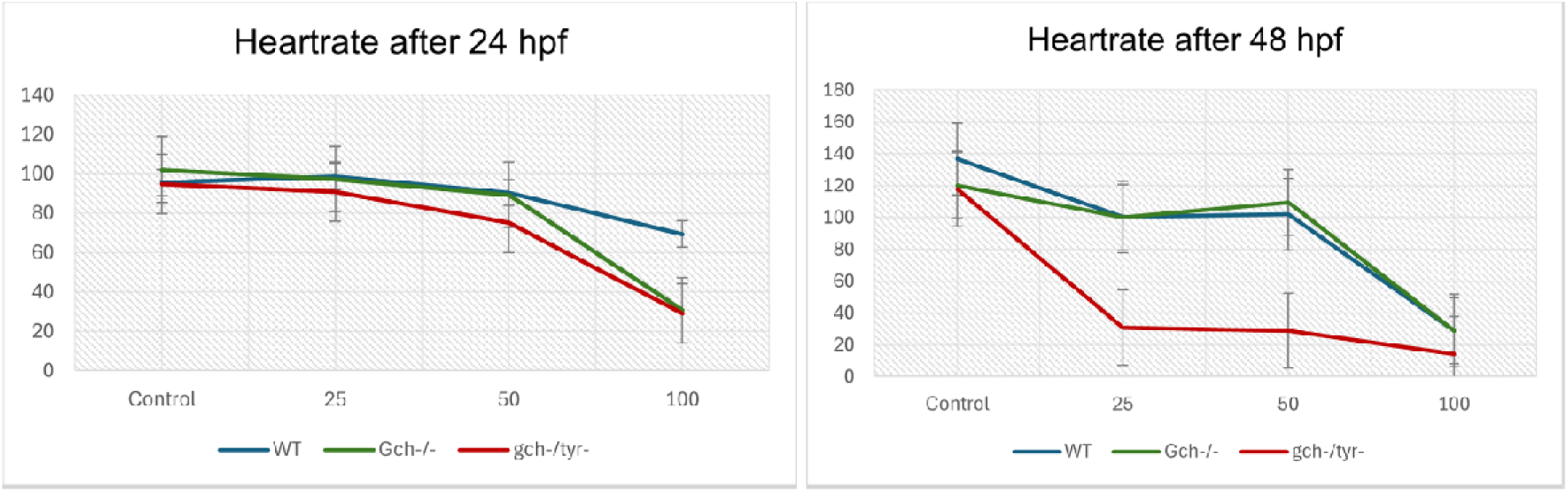
Heart rate is reduced by UV exposure depending on the genotype. Heart rate was measured after UV exposure in the WT, *gch* and *gch/tyr* embryos at 24 hpf and 48 hpf.

### UV-Induced Gene Expression

To examine molecular and genetic mechanisms of UV mediated cellular stresses and damages, we have subsequently examined gene expression using qPCR. Total RNAs were extracted from UV exposed embryos with WT, *gch* and *gch/tyr* strains. Complementary DNA was synthesised and used for examining gene expression levels of oxidative response gene *nox1*, heat response gene *hsp90*, and DNA damage repair genes *atm*, *atr* and cell cycle arrest gene *cdkn1*a. Following UV exposure, expression levels showed observable variations. Overall, elevated expression levels were observed across all genes tested after UV exposure, particularly among embryos exposed to 50 and 100 mJ/cm^2^ doses. Nevertheless, *gch^-^/^-^/tyr^-^/^-^*exhibited the highest gene expression level compared to WT and *gch^-^/^-^*. Additionally, *gch^-^/^-^* showed some elevated sensitivity to UV compared with WT (Fig. 7).

**Fig. 7.**
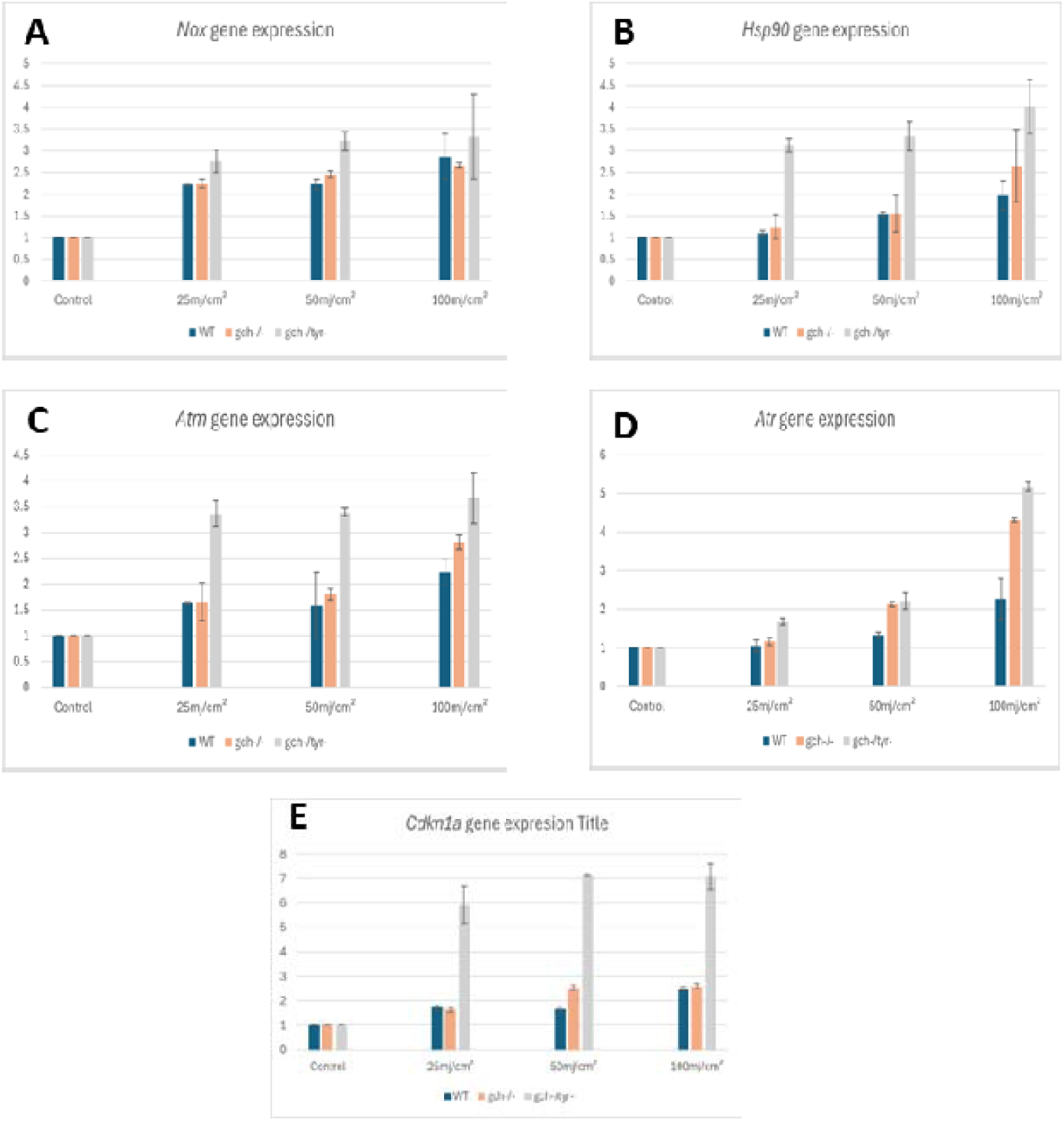
Quantitative PCR analysis of UV responding gene. Gene expression of *nox1* (A), *hsp90* (B), *atm* (C), *atr* (D) and *cdkn1a* (E). N= 3 (with 5 embryos per replicate). Values plotted are means ± SEM.

Although the overall trend is conserved, the dose responses in each gene showed some unique patterns. Firstly, *nox1* expression was upregulated by UV in a dose dependent manner in all three strains, but no significant difference between WT and mutants except a slight upregulation of the gene expression in the *gch/tyr* double mutant. In contrast, all other genes showed elevated responses in the pigment mutants against WT with varied pattern of sensitivity (Fig. 7A)

In contrast, the cell cycle arrest gene *cdkn1a* was the most prominent marker gene being highly induced in the *gch/tyr* double mutant from the lowest dose of UV, 25mJ/cm^2^. However, this gene was not highly induced by UV in the WT and *gch* mutant (Fig. 7E).

The three remaining genes, *hsp90, atm* and *atr* exhibited the intermediate responses compared to the two genes mentioned above, suggesting that their expression is influenced by dose and both pigments in a synergistic manner (Fig. 7B-D).

## Discussion

The combination of different pigment cells and their arrangement in the skin determine the overall coloration of a fish, playing a key role in camouflage, attracting mates, and sending warning signals. Pigment cells also absorb and reflect light of different wavelengths, play a crucial role in thermoregulation and protection from UV. In this study we describe the generation of *gch* mutant and *gch/tyr* double mutant in the Arabian killifish using the CRISPR/Cas9 genome editing. Our results represent the successful mutation estimated by the loss of fluorescent pigment and melanin pigment with about 88% and 83% respectively in the first generation. This suggests our strategy to knock out a gene by mixing two crRNAs would effectively generate gene knock out mutation in the Arabian killifish genome.

Our data demonstrated how pigment cells containing melanin and pteridine are essential in UV protection in fish embryos, and cells without such pigments become easily damaged by UV. The common deformities in fish embryos induced by UV are curvature or twisting of the spine^25^, reduction in growth and delay of hatching that possibly results from decreased metabolic rate similar to recorded responses in adult cichlid fish (*Cichlasoma nigrofasciatum*)^29^. Also, irritated embryos showed blood accumulation in their body, which might be a response to heartbeat changes or injury of blood vessels.

Svitačová *et al.* (2023)^30^ revealed that albino fish are more sensitive to UV stress than coloured fish in aquaculture, where pigmentation protects against ultraviolet (UV) radiation by absorbing UV radiation, mitigating genotoxic stress. Research on *Xiphophorus hybrids* by Funayama *et al*. (1993)^31^ demonstrated that melanin pigmentation potentially offers protection, as darker fish exhibited reduced UV-induced DNA damage. In addition, albino olive flounders treated with UV had significantly lower survival rates than those not treated with UV ^32^. As well as melanin also acts as an antioxidant and generates free radicals when exposed to UV light^33,34^.

In our results, the pigmented cells exhibited aggregation after UV exposure in the three lines and displayed more dendritic pattern in WT *A. dispar* as compared to non-treated pigmented cells. Depending on the light intensity, aggregation of both melanophores and fluoroleucophores responses progressed^35^ and they were destroyed at high levels of UV doses.

Previous studies have shown that ultraviolet (UV) radiation in fish can cause a range of potential effects, including direct DNA damage, oxidative stress, and phototoxicity^28,36,37^. To examine such cellular toxicities, we tested expression of genes related to oxidative stresses (*nox1*), protein damage (*hsp90*), and DNA damage (*atm, atr* and *cdkn1a*). All of these genes were induced by UV exposure in a dose dependent manner with varying patterns of response. *nox1* was the most markedly induced gene from the lowest dose of exposure (25mJ/cm2). In addition, *nox1* is the only gene that exhibits a similar induction pattern in all three genetic strains tested. This can be explained by the known fact that *nox1* is induced by reactive oxygen species that is generated by UV. Since ROS production is a normal cellular process involved in cell maintenance, even at the lowest UV doses and regardless of pigment pattern, UV may induce a normal level of oxidative stress in all three genetic strains. On the other hand, other genes, such as *hsp90* is induced by the increase of denatured proteins while *atm*, *atr*, and *cdkn1a* are induced by DNA damage. This suggests that these genes are upregulated in response to cellular damage. Since pigment mutants had more damages in the cell compared to the WT, gene expression of these genes are more highly activated in the pigment mutants in a dose dependent manner. *gch* mutant exhibited slightly elevated gene expression responses compared to WT, while *gch/tyr* double mutant showed significantly higher responses in *hsp90, atm, atr* and *cdkn1a.* These findings suggest that both pteridine in the fluoroleucophore and melanin in the melanophore play important roles in protecting cells from UV.

For *nox1* and *cdkn1a*, the dose response between WT and *gch* mutant was very similar. However, in the case of *hsp90*, *atm* and *atr*, the *gch* mutant exhibited an elevated response. This might be due to the mechanisms of UV protection by pteridine fluorescent pigment. Melanin can absorb light therefore can generate heat, but pteridine emit light and does not generate heat when the cells are exposed to UV. Therefore, in the pteridine defective mutant, *gch*, heat stress can be induced which may lead to upregulation of hsp90. Besides *hsp90, atm* and *atr* showed elevated expression in the *gch* mutant compared with WT. This is also an interesting observation that suggests important role of pteridine in protecting DNA from damage. One possibility is photoreactivation by fluorescent pigment, which could repair the DNA damage caused by UV. In the *gch* mutant, photoreactivation might be reduced and DNA damage could be enhanced causing enhanced expression of *atm* and *atr*.

*cdkn1a* exhibited the highest response to the lowest dose in the *gch/tyr* double mutant (6 folds increase compared to UV untreated control). In contrast, the gene response was quite mild in the WT and *gch* mutant. These findings also suggest the mechanisms of UV protection by melanin/melanophore and pteridine/fluoroleucophore are distinct and may be related to the issue of heat or photoreactivation, as discussed above.

*cdkn1a/p21* is responsible for inhibiting cyclin-dependent kinases (CDKs), which regulate the progress of the cell cycle by phosphorylating target proteins. In addition, p21 regulates transcription, apoptosis, DNA repair, and cell motility^38^. Therefore, strong activation of *cdkn1a* at lowest dose in the *gch/tyr* double mutant highlights the importance of these pigment cells in maintaining cell cycle and other cellular activities. Without these pigments, even a low dose of UV, can supress major cellular activities, such as cell proliferation.

To conclude, we have discovered synergistic role of melanin and pteridine in UV protection in the Arabian killifish embryo. The gene response data suggest distinct mechanisms of UV protection between two pigment cells: *atm*, *atr* and *hsp90* more reliant on pteridine, while *cdkn1a* more reliant on melanin. In contrast, *nox1* is less dependent on these pigments, resulting in a similar response to UV in all three genetic strains. To fully elucidate the differential and synergistic roles of pteridine and melanin, further investigations are needed using whole transcriptome analyses and molecular, cellular and biochemical analyses including cell cycle, apoptosis and DNA damage analyses.

## Acknowledgement

MA was funded by Tabuk University in Saudi Arabia. RM and TK are funded by NC3Rs project grant (NC/X001121/1). We also thank to Aquatic Resource Facility staff for fish husbandry, and Aaron Geffery for his support.

## References

1 Maan, M. E. & Sefc, K. M. Colour variation in cichlid fish: developmental mechanisms, selective pressures and evolutionary consequences. Semin Cell Dev Biol 24, 516–528, doi:10.1016/j.semcdb.2013.05.003 (2013).

2 Miyamoto, K. Effects of body color luminance and behavioral characteristics on predation risk in salmonid fishes. Hydrobiologia 783, 249–256, doi:10.1007/s10750-015-2573-x (2016).

3 Hoglund, E., Balm, P. H. & Winberg, S. Behavioural and neuroendocrine effects of environmental background colour and social interaction in Arctic charr (Salvelinus alpinus). J Exp Biol 205, 2535–2543, doi:10.1242/jeb.205.16.2535 (2002).

4 Kelsh, R. N. Genetics and evolution of pigment patterns in fish. Pigment Cell Res 17, 326–336, doi:10.1111/j.1600-0749.2004.00174.x (2004).

5 Le Douarin, N. & Kalcheim, C. The neural crest. 2nd edn, (Cambridge University Press, 1999).

6 Parichy, D. M. & Spiewak, J. E. Origins of adult pigmentation: diversity in pigment stem cell lineages and implications for pattern evolution. Pigment Cell Melanoma Res 28, 31–50, doi:10.1111/pcmr.12332 (2015).

7 Budi, E. H., Patterson, L. B. & Parichy, D. M. Post-embryonic nerve-associated precursors to adult pigment cells: genetic requirements and dynamics of morphogenesis and differentiation. PLoS Genet 7, e1002044, doi:10.1371/journal.pgen.1002044 (2011).

8 Singh, A. P. et al. Pigment Cell Progenitors in Zebrafish Remain Multipotent through Metamorphosis. Dev Cell 38, 316–330, doi:10.1016/j.devcel.2016.06.020 (2016).

9 Fujii, R. Cytophysiology of Fish Chromatophores. Int Rev Cytol 143, 191–255 (1993).

10 Sugimoto, M. Morphological color changes in fish: regulation of pigment cell density and morphology. Microsc Res Tech 58, 496–503, doi:10.1002/jemt.10168 (2002).

11 Hirata, M., Nakamura, K., Kanemaru, T., Shibata, Y. & Kondo, S. Pigment cell organization in the hypodermis of zebrafish. Dev Dyn 227, 497–503, doi:10.1002/dvdy.10334 (2003).

12 Parichy, D. M. Evolution of pigment cells and patterns: recent insights from teleost fishes. Curr Opin Genet Dev 69, 88–96, doi:10.1016/j.gde.2021.02.006 (2021).

13 Fujii, R. The regulation of motile activity in fish chromatophores. Pigment Cell Res 13, 300–319, doi:10.1034/j.1600-0749.2000.130502.x (2000).

14 Frohnhofer, H. G., Krauss, J., Maischein, H. M. & Nusslein-Volhard, C. Iridophores and their interactions with other chromatophores are required for stripe formation in zebrafish. Development 140, 2997–3007, doi:10.1242/dev.096719 (2013).

15 Kimura, T. et al. Leucophores are similar to xanthophores in their specification and differentiation processes in medaka. P Natl Acad Sci USA 111, 7343–7348, doi:10.1073/pnas.1311254111 (2014).

16 Hamied, A. et al. Identification and Characterization of Highly Fluorescent Pigment Cells in Embryos of the Arabian Killifish (Aphanius Dispar). iScience 23, 101674, doi:10.1016/j.isci.2020.101674 (2020).

17 Frenkel, V. & Goren, M. Factors affecting growth of killifish, Aphanius dispar, a potential biological control of mosquitoes. Aquaculture 184, 255–265, 10.1016/S0044-8486(99)00326-9 (2000).

18 Esmaeili, H. R., Asrar, T. & Gholamifard, A. Cyprinodontid fishes of the world: an updated list of taxonomy, distribution and conservation status (Teleostei: Cyprinodontoidea). Iranian Journal of Ichthyology 5, 1–29 (2018).

19 Plaut, I. Resting metabolic rate, critical swimming speed, and routine activity of the euryhaline cyprinodontid, Aphanius dispar, acclimated to a wide range of salinities. Physiol Biochem Zool 73, 590–596, doi:10.1086/317746 (2000).

20 Alsakran, A. et al. Stage-by-stage exploration of normal embryonic development in the Arabian killifish, Aphanius dispar. Dev Dyn, doi:10.1002/dvdy.738 (2024).

21 Madronich, S., McKenzie, R. L., Björn, L. O. & Caldwell, M. M. Changes in biologically active ultraviolet radiation reaching the Earth’s surface. J Photoch Photobio B 46, 5–19, doi:Doi 10.1016/S1011-1344(98)00182-1 (1998).

22 McKenzie, R. L., Aucamp, P. J., Bais, A. F., Bjorn, L. O. & Ilyas, M. Changes in biologically-active ultraviolet radiation reaching the Earth’s surface. Photochem Photobiol Sci 6, 218–231, doi:10.1039/b700017k (2007).

23 El-Nouby, A. M. Effect of stratospheric ozone in UVB solar radiation reaching the Earth’s surface at Qena, Egypt. Atmospheric pollution research 1, 155–160 (2010).

24 Diaz, S. et al. Ozone and UV radiation over southern South America: climatology and anomalies. Photochem Photobiol 82, 834–843, doi:10.1562/2005-09-26-RA-697 (2006).

25 Dong, Q., Svoboda, K., Tiersch, T. R. & Monroe, W. T. Photobiological effects of UVA and UVB light in zebrafish embryos: evidence for a competent photorepair system. J Photochem Photobiol B 88, 137–146, doi:10.1016/j.jphotobiol.2007.07.002 (2007).

26 Mahmoud, U. M., Mekkawy, I. A. & Sayed Ael, D. Ultraviolet radiation-A (366 nm) induced morphological and histological malformations during embryogenesis of Clarias gariepinus (Burchell, 1822). J Photochem Photobiol B 95, 117–128, doi:10.1016/j.jphotobiol.2009.02.003 (2009).

27 Torres Nunez, E., et al. Molecular response to ultraviolet radiation exposure in fish embryos: implications for survival and morphological development. Photochem Photobiol 88, 701–707, doi:10.1111/j.1751-1097.2012.01088.x (2012).

28 Zang, L., Shimada, Y., Miyake, H. & Nishimura, N. Transcriptome analysis of molecular response to UVC irradiation in zebrafish embryos. Ecotoxicol Environ Saf 231, 113211, doi:10.1016/j.ecoenv.2022.113211 (2022).

29 Winckler, K. & Fidhiany, L. Significant influence of UVA on the general metabolism in the growing Cichlid fish, Cichlasoma nigrofasciatum. Journal of Photochemistry and Photobiology B: Biology 33, 131–135, 10.1016/1011-1344(95)07238-1 (1996).

30 Svitačová, K., Slavík, O. & Horký, P. Pigmentation potentially influences fish welfare in aquaculture. Applied Animal Behaviour Science 262, 105903, 10.1016/j.applanim.2023.105903 (2023).

31 Funayama, T., Mitani, H. & Shima, A. ULTRAVIOLET-INDUCED DNA DAMAGE AND ITS PHOTOREPAIR IN TAIL FIN CELLS OF THE MEDAKA, Oryzias lutipes. Photochemistry and photobiology 58, 380–385 (1993).

32 Fukunishi, Y. et al. Comparison of UV-B tolerance between wild-type and albino Japanese flounder Paralichthys olivaceus juveniles. Aquaculture Science 65, 149–152 (2017).

33 Aspengren, S., Hedberg, D., Sköld, H. N. & Wallin, M. New insights into melanosome transport in vertebrate pigment cells. International review of cell and molecular biology 272, 245–302 (2008).

34 Nilsson Skold, H., Aspengren, S. & Wallin, M. Rapid color change in fish and amphibians - function, regulation, and emerging applications. Pigment Cell Melanoma Res 26, 29–38, doi:10.1111/pcmr.12040 (2013).

35 Arpigny, J. et al. Factors influencing motile activities of fish chromatophores. Advances in Comparative and Environmental Physiology: Volume 20, 1–54 (1994).

36 Sayed, A. E.-D. H. & Mitani, H. Immunostaining of UVA-induced DNA damage in erythrocytes of medaka (Oryzias latipes). Journal of Photochemistry and Photobiology B: Biology 171, 90–95 (2017).

37 Wagner, J. T. & Podrabsky, J. E. Extreme tolerance and developmental buffering of UV-C induced DNA damage in embryos of the annual killifish Austrofundulus limnaeus. J Exp Zool A Ecol Genet Physiol 323, 10–30, doi:10.1002/jez.1890 (2015).

38 Dutto, I., Tillhon, M., Cazzalini, O., Stivala, L. A. & Prosperi, E. Biology of the cell cycle inhibitor p21(CDKN1A): molecular mechanisms and relevance in chemical toxicology. Arch Toxicol 89, 155–178, doi:10.1007/s00204-014-1430-4 (2015).

